# THUNDER: A reference-free deconvolution method to infer cell type proportions from bulk Hi-C data

**DOI:** 10.1101/2020.11.12.379941

**Authors:** Bryce Rowland, Ruth Huh, Zoe Hou, Jia Wen, Yin Shen, Ming Hu, Paola Giusti-Rodríguez, Patrick F Sullivan, Yun Li

## Abstract

Hi-C data provide population averaged estimates of three-dimensional chromatin contacts across cell types and states in bulk samples. Effective analysis of Hi-C data entails controlling for the potential confounding factor of differential cell type proportions across heterogeneous bulk samples. We propose a novel unsupervised deconvolution method for inferring cell type composition from bulk Hi-C data, the Two-step Hi-c UNsupervised DEconvolution appRoach (THUNDER). We conducted extensive simulations to test THUNDER based on combining two published single-cell Hi-C (scHi-C) datasets. THUNDER more accurately estimates the underlying cell type proportions compared to supervised and unsupervised methods (e.g., MuSiC, TOAST, and NMF). We further demonstrate the practical utility of THUNDER to estimate cell type proportions and identify cell-type-specific interactions in Hi-C data from adult human cortex tissue samples. THUNDER will be a useful tool in adjusting for varying cell type composition in population samples, facilitating valid and more powerful downstream analysis such as differential chromatin organization studies. Additionally, THUNDER estimated contact profiles provide a useful exploratory framework to investigate cell-type-specificity of the chromatin interactome while experimental data is still rare.

## Introduction

Statistical deconvolution methods have been applied extensively to studies of gene expression and DNA methylation to infer cell type proportions and estimate cell-type-specific profiles(1–6). Deconvolution methods infer latent clusters from observed data which can correspond to either cell types or cell states (hereafter we refer to both as cell types). In epigenome-wide association studies (EWAS) where the individual-level signal is a mixture of methylation profiles from different cell types, it has become standard practice to control for inferred cell type proportions when analyzing heterogeneous samples.(7) As we accumulate chromatin interaction information from heterogeneous samples using recently developed technologies such as Hi-C at an increasing rate, there will soon be sufficient individual level data to conduct similar 3D-chromatin-interactome wide association studies (3WAS) or chromatin interactome QTL (iQTL) studies.(8) Similar to DNA methylation and gene expression, there is growing evidence from single-cell Hi-C (scHi-C) data of important cell-to-cell variability in spatial chromatin interaction.(9–12) In order to effectively garner insights from associations between chromatin interactions and phenotypes of interest or to identify genetic determinants underlying variations in 3D-chromatin-interactome across biological samples, future 3WAS or iQTL analyses must control for the almost inevitable confounding factor of differential cell type proportions across heterogeneous bulk samples. If not accounted for, we risk inducing an increased false positive rate by Simpson’s Paradox.(7,13) However, to the best of our knowledge, there is no statistical deconvolution method which is capable of leveraging both intrachromosomal and interchromosomal contacts for deconvolution across multiple bulk Hi-C samples simultaneously.

There exist two particular challenges of performing deconvolution in bulk Hi-C data: a lack of cell-type-specific Hi-C reference profiles and having no ubiquitous aggregating unit for summarizing Hi-C data. First, many deconvolution methods require cell-type-specific reference profiles for each cell type potentially present in a mixture (e.g., the genes whose expression define a cell type), but analogous data are not yet available for Hi-C. Second, Hi-C data can be summarized at several different structural levels, such as A/B compartments, topologically associating domains (TADs)(14), frequently interacting regions (FIREs)(15,16), chromatin loops(17), interchromosomal contacts, and/or intrachromosomal contacts(18,19), and it is unclear which level(s) of measurement are most scientifically relevant or effective for deconvolution purposes. In contrast, when deconvolving gene expression data it is clear that the aggregating unit of interest is the gene.

To address these challenges, we propose a non-negative matrix factorization (NMF) based Two-step Hi-c UNsupervised DEconvolution appRoach (THUNDER), to infer cell type proportions from bulk Hi-C data. THUNDER consists of a feature selection step and a deconvolution step, both of which rely on NMF. NMF has been used in many computational biology applications to cluster genes, discover cancer types using microarray data, and study functional relationships of genes(20–22). In the first step, we perform feature selection on the cell type profiles estimated from an initial deconvolution to identify informative bin-pairs in the mixture data (Figure 1a,b). In the second step, we perform deconvolution after subsetting the mixture matrix on informative bin-pairs (Figure 1c).

**Figure 1.**
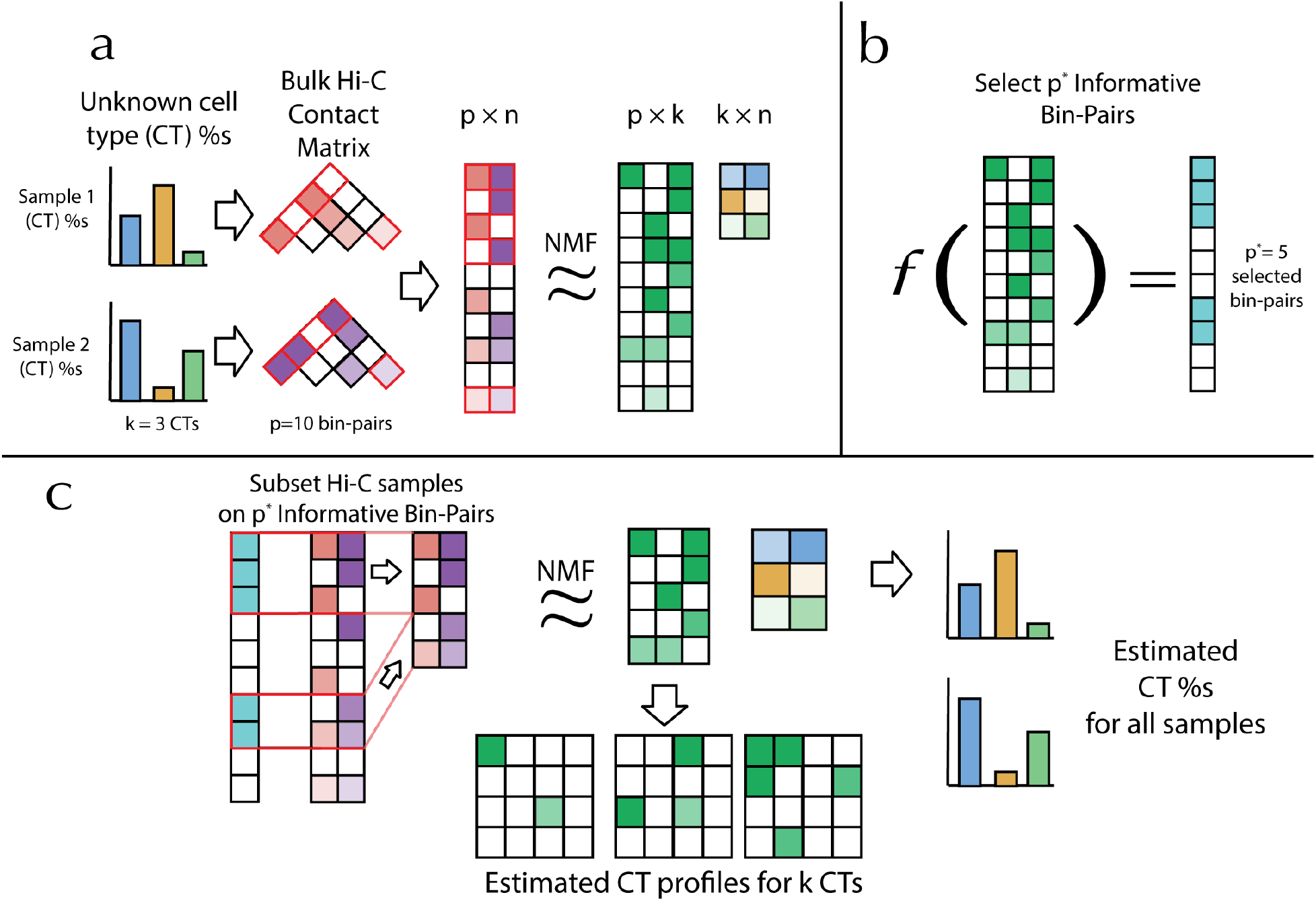
Overview of *THUNDER* Procedure. (a) Overview of nonnegative matrix factorization (NMF) in the context of bulk Hi-C data. Three underlying cell types each contribute to the observed contact frequencies in two bulk Hi-C samples. The first step of the *THUNDER* algorithm is to deconvolve the input bulk Hi-C data into two estimated matrices: the cell type profile matrix and the proportion matrix. (b) In order to select informative bin-pairs for deconvolution, *THUNDER* utilizes a feature selection algorithm specifically tailored to Hi-C data to analyze the contact frequency distribution of the bin-pairs in the cell type profile matrix. (c) In the final step of *THUNDER*, we subset the bin-pairs contained in the input bulk Hi-C samples to only informative bin-pairs and perform NMF a second time. The proportion matrix is scaled to be estimates of the underlying cell type proportions in the bulk Hi-C samples. The cell type profile matrix estimates cell-type-specific contact distributions.

To the best of our knowledge, THUNDER is the first unsupervised deconvolution method for Hi-C data that integrates both intrachromosomal and interchromosomal contact information to estimate cell type proportions in multiple bulk Hi-C samples simultaneously. Two other matrix-based deconvolution approaches exist for Hi-C intrachromosomal contact matrices: 3CDE infers non-overlapping domains of chromatin activity in each cell type and uses a linear combination of binary interaction information at these domains to perform deconvolution.(23) Junier *et al.* put forth a method to infer overlapping domains of chromatin activity as well as their mixture proportions.(24) Unlike THUNDER, neither method integrates information from interchromosomal bin-pairs. We tested 3CDE on our simulated bulk Hi-C mixtures, but found that it is almost impossible to apply in practice because it does not accommodate the inclusion of interchromosomal contacts and it requires across-sample cell type matching to align proportion estimates since it infers cell type proportions for each sample separately (Supplementary Figure 1). To the best of our knowledge, no software accompanies the work by Junier *et al.(24)* Carstens *et al.* infer chromatin structure ensembles from bulk Hi-C contact information using a Bayesian approach but does not infer cell type proportions directly.(25)

In this work, we consider two other deconvolution methods developed for gene expression or methylation data: MuSiC and TOAST. MuSiC is a reference based deconvolution method which estimates cell type proportions from bulk RNA sequencing data based on multi-subject single cell data.(6) TOAST is a feature selection algorithm for gene expression or methylation data to select a pre-specified number of features while performing unsupervised deconvolution via NMF.(3)

## Results

### THUNDER Feature Selection

In order to determine the feature selection method for THUNDER, using scHi-C data generated from Ramani *et al.* (10), we simulated 12 mixtures of Hi-C data at 10Mb resolution consisting of three cell lines, HAP1, HeLa, and GM12878, where we set the cell composition proportions (details in **Methods**). We evaluated the performance of 11 published and novel NMF feature selection strategies for intrachromosomal only and interchromosomal only bin-pairs (see **Supplementary Table 1** for definitions).

Our simulation results suggest that the optimal feature selection method differs for deconvolving interchromosomal and intrachromosomal contacts (Figure 2). For intrachromosomal contacts, the best feature selection method is “High CTS (median)” which prioritizes features with high cell-type-specificity using median-based empirical thresholds and selects an average of 353 informative bin-pairs out of an average of 2,590 input intrachromosomal contact features. The best performing interchromosomal feature selection method is “High ACV”. “High ACV” prioritizes features with high across-cell-type variation (ACV) using mean-based empirical thresholds and selects an average of 287 informative bin-pairs out of an average of 42,871 input interchromosomal contact features. We refer to these two methods hereforeward as THUNDER-intra and THUNDER-inter, respectively. Compared to NMF with no feature selection, THUNDER-intra reduced average MAD (mean absolute deviation, smaller indicates better performance) by 42% and increased average Pearson correlation by 0.4%. Similarly, THUNDER-inter reduced average MAD by 69% and increased average Pearson correlation by 3.3%. Feature selection methods that require specifying the number of informative bin-pairs *a priori* such as Fano-100 and Fano-1000, which selects the top 100 and 1000 features with highest Fano factor respectively, exhibit the most variable performance across simulations, and perform poorly relative to other methods despite specifying a similar number of bins.

**Figure 2.**
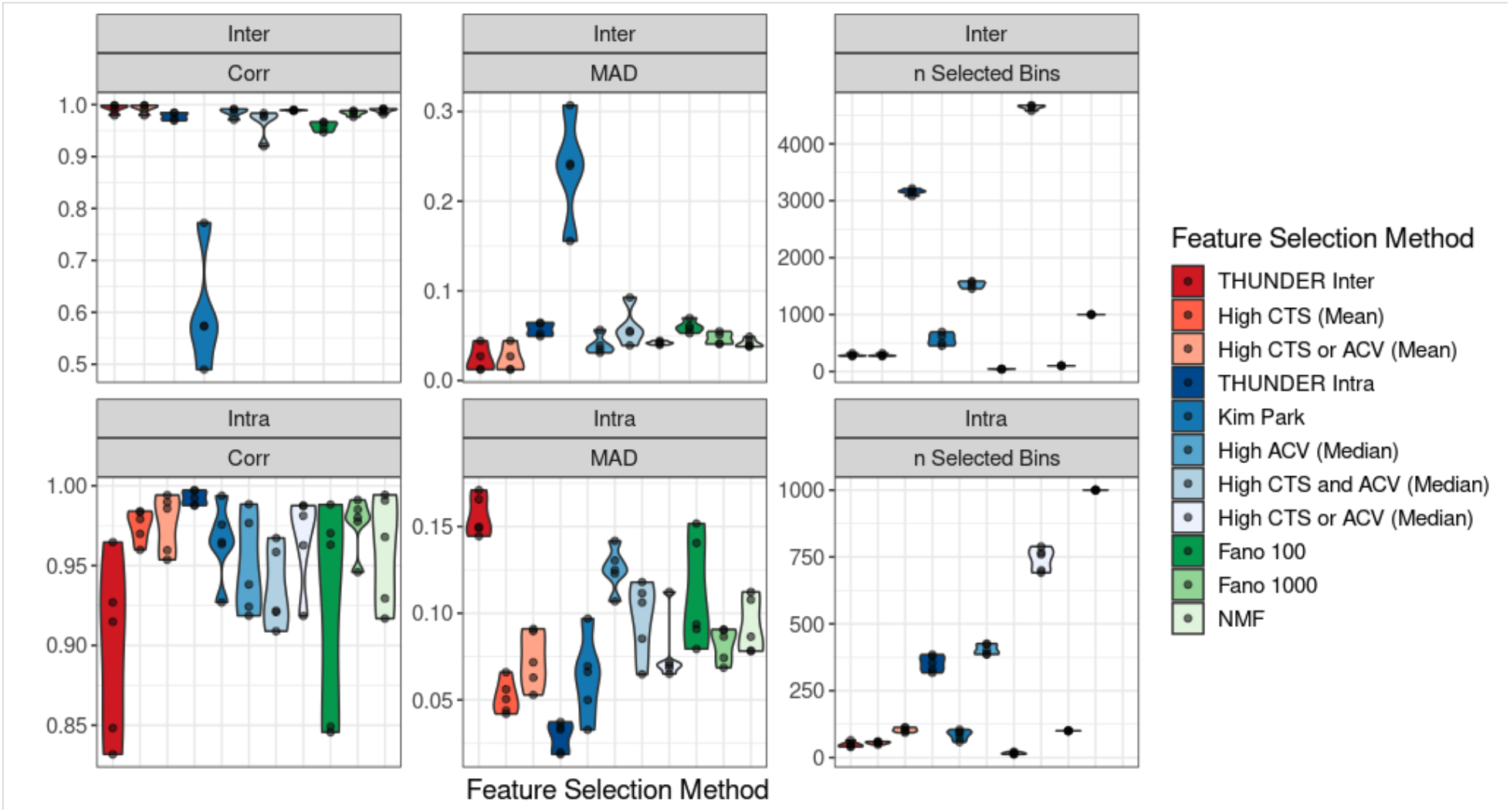
Performance of Feature Selection Strategies for Unsupervised Hi-C Deconvolution in HAP1, HeLa, and GM12878 Mixtures. We test 11 feature selection strategies including no feature selection (NMF), Fano 100, Fano 1,000, and 8 feature selection strategies combining bin-pairs with high cell-type-specificity (CTS) and high across-cell-type variation (ACV). Colors are grouped such that the “reds” are strategies analyzing the estimated cell-type-specific profiles using the mean across bin-pairs for thresholding, “blues” are feature score strategies analyzing the estimated cell-type-specific profiles using the median across bin-pairs for thresholding, and “greens” are NMF with no feature selection or a pre-specified number of features based on Fano factor. Distributions are presented across simulation replicates.

### Simulations based on scHi-C from brain (Lee et al)

We tested the accuracy of THUNDER cell type proportion estimates using scHi-C data from Lee *et al*.(12) to simulate 18 Hi-C mixtures at 10Mb resolution of 6 brain cell types: microglia, astrocytes, oligodendrocytes, oligodendrocyte progenitor cells, endothelial cells, and neuronal cells. THUNDER cell type proportion estimates were most accurate when deconvolving intrachromosomal and interchromosomal contacts together, reducing MAD by 9.7% and 7.6% and increasing Pearson correlation by 3.7% and 1.4% compared to intrachromosomal contacts and interchromosomal contacts respectively. We compared THUNDER’s performance to NMF with no feature selection, MuSiC, and TOAST on mixtures with both intrachromosomal and interchromosomal contacts, intrachromosomal contacts only, and interchromosomal contacts only (Figure 3). THUNDER outperformed all alternative reference-free deconvolution approaches in each simulation. When deconvolving both intrachromosomal and interchromosomal contacts together, THUNDER decreased average MAD by 23% and 36% and increased Pearson correlation by 6% and 31% relative to NMF and TOAST, respectively. MuSiC, a reference-based deconvolution approach, outperformed THUNDER in all simulation scenarios when all cell types in the mixtures are present in the reference panel. However, due to the current paucity of cell-type specific Hi-C reference panels, we tested the performance of MuSiC with one and two cell types randomly removed from the reference panel (**Methods**). In all three simulation settings, MuSiC’s performance decreased with the number of cell types randomly removed from the reference (MuSiC, MuSiC - One Missing, and Music - Two Missing in Figure 3a,b). The performance of MuSiC one-missing was comparable to THUNDER in all simulation settings, and MuSiC - Two Missing was either worst or close to the worst performing methods. From our simulations, THUNDER performed best among reference free methods, and was more robust compared to MuSiC which performed poorly when cell types are missing from the reference panel. We anticipate reference based methods such as MuSiC will become more advantageous as we accumulate resources to build a comprehensive reference panel. Currently, with limited resources to construct a reference dataset, reference free methods are more valuable.

**Figure 3.**
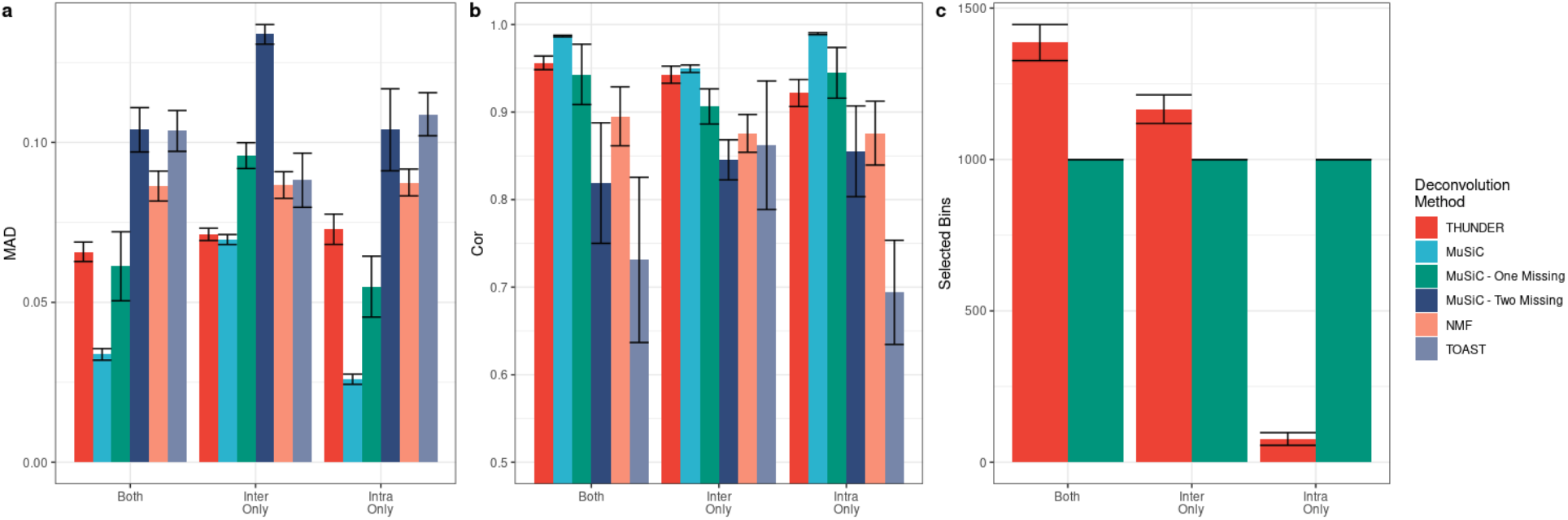
Performance of Deconvolution Methods on Mixtures with 6 Human Brain Cell Types. (a,b) The average mean absolute deviation (MAD) and average Pearson correlation comparing the true underlying cell type proportions to the simulated true proportions across simulations across 5 simulation replicates. Lower MAD and higher Pearson correlation indicates better performance. Error bars are equal to the standard deviation across simulation replications. (c) Number of bin-pairs selected by deconvolution methods which perform feature selection.

**Figure 4.**
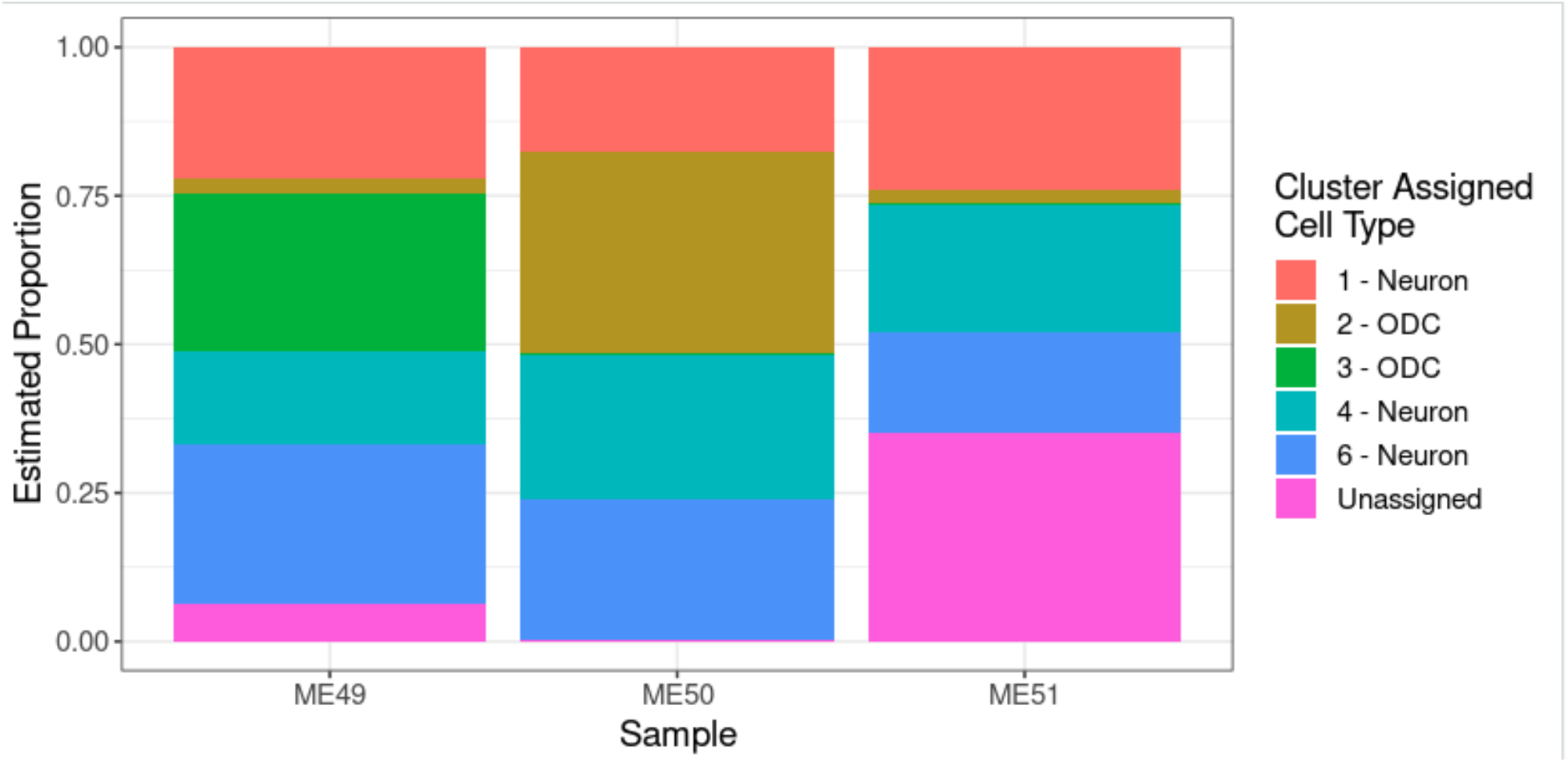
THUNDER Estimated Cell Type Proportions in 3 Samples of Human Cortex Tissue. We use THUNDER to estimate cell type proportions for 3 Hi-C samples from cortex tissue and perform enrichment analyses to assign brain cell types to THUNDER clusters. Our results match the expected ratio of neuronal to non-neuronal cells in cortex tissue.

### Simulations based on scHi-C from brain (Giusti-Rodriguez et al.)

We applied THUNDER to bulk Hi-C data generated on cortex tissue from three postmortem adult samples (**Methods**). In downstream analysis, we proceeded with the deconvolution results when k=6 due to the greatest consistency across samples (see Supplemental Figure 2).

In order to assign plausible cell type labels to the 6 THUNDER inferred clusters, we compared the cluster-specific bins to cell-type specific enhancers and genes from four cell types commonly found in cortex tissue. 5 out of 6 THUNDER features (all except THUNDER cluster 4) demonstrated enrichment for neuronal enhancers (p < 0.05/48 = 1.04e-3), so we assigned each cluster to a cortical cell type based on other significant enrichments if possible. THUNDER cluster 1 showed evidence of enrichment for neuronal specifically expressed genes (p = 3.2e-3) and was thus assigned as neurons. THUNDER cluster 6 demonstrated enrichment for neuronal enhancers (p = 3.79e-9) and a trend (although not statistically significant) for enrichment of neuron specific genes (p = 0.104). We assigned THUNDER cluster 6 to neurons. THUNDER cluster 4 demonstrated enrichment with neuronal enhancers (p = 1.89e-3), and was thus assigned to neurons as well. Bins distinct to THUNDER clusters 2 and 3 demonstrated consistent evidence of enrichment of oligodendrocytes (ODC) features, in terms of enhancers (p = 3.3e-4 and p = 7.5e-9) and ODC-specifically expressed genes (p = 7.5e-3 and p = 4.78e-3). Therefore, both were assigned as ODC cells. THUNDER cluster 5 was not assigned to a cell type due to a lack of specific enrichments.

With these assigned cell type labels to the clusters, THUNDER estimated 62.7-65.2% neurons, 2.3-34.5% ODCs, and 0.3-35% unassigned for the three samples, largely matching the expected ratio of neuronal to non-neuronal cells in cortex tissue. Additionally, THUNDER informative bin-pairs identified biologically relevant cell-type specific interactions. For example, the bin-pair defined by genomic regions chr5:130Mb-131Mb and chr5:131Mb-132Mb was an informative bin pair for THUNDER cluster 6, which was assigned to neurons via enrichment analysis. This bin-pair contained 14 high-confidence regulatory chromatin interactions (HCRCI) identified in the three adult cortical samples in a previous study with genomic coordinates within chr5:130600000-130970000 and chr5:131100000-131730000, respectively.(26) Further, two neuron-specific genes identified in our analysis of data from Zhang *et al.* were contained in chr5:131,100,000-131,730,000, *ACSL6* and *P4HA2*. Together, these results suggest that this THUNDER informative bin pair may correspond to a group of neuron-specific chromatin interactions. Another such example is the THUNDER informative bin-pair defined by the genomic regions chr12:121Mb-122Mb and chr12:122Mb-123Mb for THUNDER cluster 3, which enrichment analysis suggested as ODCs. The two regions defining this bin pair contained 64 HCRCIs, and two ODC specifically expressed genes, *P2RX7* and *ANAPC5*. Our results suggest that THUNDER estimated cell-type-specific profiles (see Supplemental Table 1) can identify biologically meaningful cell-type-specific interactions from bulk Hi-C data.

### Computations on 10Kb Hi-C data

THUNDER scales linearly with both the number of samples under inference and the number of input features (Supplementary Tables 1-3). We assessed THUNDER’s computing performance on Hi-C data of lymphoblastoid cell lines (LCLs) derived from five YRI (Yoruba in Ibadan, Nigeria) individuals.^7^ Specifically, we analyzed intrachromosomal contacts at 10Kb resolution, with 38,343,298 unique intrachromosomal bin-pairs ranging from 380,000 to 3.5 million bin-pairs per chromosome. To obtain cell type proportion estimates genome-wide using THUNDER, we first perform feature selection by chromosome, then concatenate the selected features across chromosomes as input for the final deconvolution estimate. THUNDER’s average computing time is 3.4 hours (range 0.6-7.2 hours) with an average of 57GB memory (range 18GB - 103GB) per chromosome using a single core on a 2.50 GHz Intel processor with 256GB of RAM. The final genome-wide estimation step to obtain cell type proportions, with 693,771 (~2%) bin-pairs selected as informative, took 2.5 hours and 18GB of memory (Supplementary Table 2). Similar summaries are presented for analyzing 3 and 10 YRI samples respectively (Supplementary Tables 1 and 3). One advantage of THUNDER’s feature selection method when analyzing genome-wide Hi-C data is the ease with which it can be parallelized by subsetting the original input matrix in smaller regions than by chromosome, then concatenating Hi-C data for the final cell type proportion estimation step. This run time and memory usage serves as an upper limit on the computational costs of running THUNDER, as 10Kb is one of the finest resolutions of Hi-C data currently analyzed in practice.

## Discussion

THUNDER is the first unsupervised deconvolution method for Hi-C data that integrates both intrachromosomal and interchromosomal contact information to estimate cell type proportions in multiple bulk Hi-C samples. Across all simulations, THUNDER’s accuracy in estimating cell type proportions exceeded all reference-free alternative approaches tested. Importantly, THUNDER’s feature selection strategy for identifying informative bin-pairs before deconvolution improves performance relative to NMF with no feature selection. We found THUNDER to be a robust alternative to reference-dependent methods which may not estimate cell type proportions accurately when cells are missing from the reference panel, a realistic scenario in practice with Hi-C data deconvolution. Further, we found that even in non-cancerous cell lines, the inclusion of sparse interchromosomal contact information (in addition to intrachromosomal contacts) improves deconvolution performance. This, however, comes at the cost of increased computational cost. THUNDER also provides an approach to infer cell-type-specific contact frequency from bulk Hi-C data.

We demonstrated that THUNDER successfully integrates interchromosomal contacts to improve deconvolution estimates for Hi-C data. In most cell types, we have more reliable Hi-C data at a much larger number of intrachromosomal bin-pairs compared to interchromosomal bin-pairs. For this reason, previous methods to deconvolve Hi-C data restricted their estimation to these intrachromosomal contacts. However, even in simulations with no strong interchromosomal signatures (for example, in the Lee et al human brain data), THUNDER’s performance improves when integrating interchromosomal and intrachromosomal data for deconvolution relative to only using intrachromosomal contacts. Our results suggest some value in including interchromosomal contacts bulk Hi-C deconvolution, though at the tradeoff of computational efficiency. Since we analyze Hi-C data by grouping contacts into bin-pairs, the feature space increases rapidly with increasing bins. As demonstrated in our computation test, THUNDER’s computation costs increase linearly as the number of features increases. Despite this tradeoff, our results suggest that interchromosomal bin-pairs contain useful information that warrant consideration before excluding these bin-pairs in Hi-C deconvolution.

Additionally, we demonstrate that THUNDER estimated cell-type-specific profiles are enriched for relevant cell-type-specific enhancers and specifically expressed genes through our analysis of 3 adult human cortex samples. We demonstrate how existing cell-type specific annotations can be used to label THUNDER inferred clusters, and thus provide cell type proportion estimates in real Hi-C data. Thus, the estimated cell type profile matrix serves a dual purpose: identifying informative bin-pairs from the large input feature space (dimension reduction) and accurately estimating relative cell-type-specific contact frequency at informative bin-pairs.

An additional application of these cell-type-specific contact profiles could be in fine mapping of GWAS variants in non-coding regions of the genome. Genome-wide association studies (GWAS) have identified over 200,000 unique associations between single-nucleotide polymorphisms (SNPs) and common diseases or traits of interest.(27) However, the majority of these SNPs reside in non-coding regions where little is understood about their underlying functional mechanisms, which has limited the adoption of variant-trait associations into revealing molecular mechanisms and further into transforming clinical practice. Functional annotation of GWAS results are often most relevant in a cell-type-specific fashion due to important variability across cell types(28). By further understanding the cell-type-specific interactome via THUNDER’s estimated profiles, we anticipate more informative linking putatively causal variants identified by GWAS to the target genes on which they act.

While we have presented results for Hi-C data here, the THUNDER algorithm could easily be modified to other variations of Hi-C data such as HiChIP/PLAC-seq data (HP data), which couple standard Hi-C with chromatin immunoprecipitation to profile chromatin interactions anchored at genomic regions bound by specific proteins or histone modifications, with reduced cost and enhanced resolution.(29,30) Used in concert with methods to identify long-range chromatin interactions from HP data(31), our method is anticipated to efficiently leverage interchromosomal contacts jointly with high quality intrachromosomal contacts to estimate underlying cell type proportions. The robustness of our feature selection strategy and subsequent deconvolution performance warrant future interrogation in the setting of HP data.

There are three primary limitations of our study. First, due to the number of cells present in current scHi-C datasets and the library size, our simulation analysis was limited to a coarse resolution of 10Mb bins when generating our synthetic bulk Hi-C data. However, we find that THUNDER still performs exceedingly well in estimating true cell type proportions even at coarse resolution. Secondly, the number of cell types and the overall coverage of the genome with our synthetic bulk Hi-C data are both much lower than one would expect in a realistic sample of bulk Hi-C data. As more scHi-C data becomes available, we hope to continue to test THUNDER in different real-data based scenarios which may be more realistic in terms of Hi-C data’s read-depth.

## Conclusion

To summarize, we present THUNDER, an unsupervised deconvolution approach tailored to the unique challenges of deconvolving Hi-C data. THUNDER accurately estimates cell type proportions in bulk Hi-C data. THUNDER’s biologically motivated feature selection approach performs well in all of our real data or real-data based simulations, including human cell lines, human cortex tissue, and human brain cells. We have demonstrated the computational efficiency of the method through our analysis of 10Kb resolution Hi-C data. Finally, the estimated cell-type-specific chromatin interactome profiles are valuable for identifying bin-pairs which interact differentially across cell types.

Accurately estimating underlying cell type proportions via THUNDER should be the first step in any individual-level differential analysis of bulk Hi-C data to control for the almost inevitable confounding factor of underlying cell type proportions. Additionally, THUNDER provides a unique tool to identify differentially interacting bin-pairs at the cell-type-specific level which can be associated with disease or phenotypes of interest. An R package for running THUNDER can be downloaded from https://github.com/brycerowland/thundeR.git. We anticipate THUNDER to become a convenient and essential tool in future multi-sample Hi-C data analysis.

## Methods

### THUNDER

In order to estimate the underlying cell type proportions found in bulk Hi-C datasets, we propose a Two Step Hi-C UNsupervised DEconvolution appRoach (THUNDER). THUNDER consists of a feature selection step and a deconvolution step, both of which rely on non-negative matrix factorization. For Hi-C data, *V*denotes the *p x n* mixture matrix of bulk Hi-C samples with *p* bin-pairs and *n* columns of mixture samples. We let *k* > 0 be an integer specified for the number of distinct cell types in the mixture sample and is chosen *a priori*. NMF seeks to find an approximation *V* ≈ *WH*, where *W* and *H* are *p* × *k* and *k* × *n* non-negative matrices. We refer to *W* and *H* as the cell type profile and proportion matrices, respectively. The NMF problem can be solved by finding a local minimum for the Euclidean norm between *V* and *WH*, || *V* − *WH* ∥^2^, under the constraint that *W* and *H* are non-negative. We use the NMF R package(32) with the updates provided by Lee and Seung(33) with random initialization of the *W* and *H* matrices.

In step one of THUNDER, we perform an initial NMF deconvolution estimate on the *p x n* matrix *V* to obtain the deconvolution estimate *V* ≈ *W*_1_*H*_1_ where *W*_1_ is a *p x k* matrix and *H*_1_ is a *k x n* matrix. We then perform feature selection using the decomposition to identify informative bin-pairs across cell types. THUNDER performs feature selection on intrachromosomal and interchromosomal contacts separately. Let *W*_1_(*i, j*) denote the element in the *i^th^* row and *j^th^* column of the cell-type-specific profile matrix *W_1_*. Let *S_intra_* and *S_inter_* denote the set of intrachromosomal and interchromosomal bin-pairs respectively.

Standard deviation across cell types for bin-pair *i* is defined as,

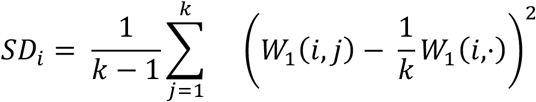

Feature score across cell types for bin-pair *i* is defined as follows.

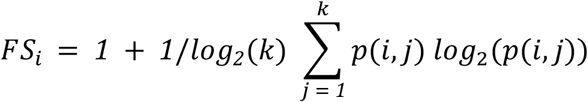

where *p*(*i*, *Ω*) is the probability that the *i*-th pairwise bin contributes to cell type Ω, i.e. 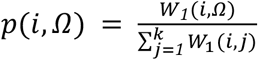. Feature scores range from [0,1] with higher scores representing bin-pairs with higher cell-type-specificity. We further define,

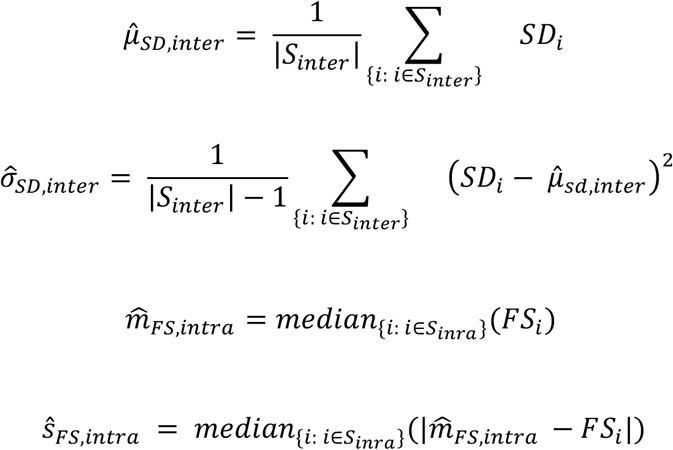

Intrachromosomal bin-pair *i* is defined to be an informative bin-pair if 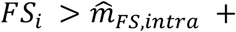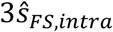, and interchromosomal bin pair j is defined to be an informative bin pair if 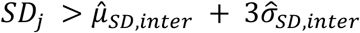.

After identifying *p*\* informative bin-pairs, we subset *V* on all informative bin-pairs to form the reduced *p*\**x n* mixture matrix *V*\*. We then perform NMF on *V*\* to arrive at our final estimates, *W*\* (*of dimension p*\* *x k*) and *H*\* (dimension *k x n*). Finally, we adjust the columns of *H*\* to sum to one to represent cell type proportions. The scaled elements of *H*\*are cell type proportion estimates in the *p* mixture samples. The columns of *W*\* are parsimonious cell-type-specific contact profiles. These parsimonious contact profiles estimate Hi-C contact frequencies at the bin-pairs which most differentiate the inferred cell types in the Hi-C samples.

### MuSiC

MuSiC is a reference based deconvolution method which estimates cell type proportions from bulk RNA sequencing data based on multi-subject single cell RNA sequencing data. MuSiC leverages features which demonstrate cross-cell and cross-sample consistency to apply cell-type-specific feature information in estimating cell type proportions. MuSiC additionally applies a tree-based procedure to address collinearity in closely related cell types within a bulk tissue. To run MuSiC, we used the MuSiC R package (version 0.1.1) with default parameters. We constructed a scHi-C reference dataset using cells from Lee et al. which match cells considered in the simulated mixtures. Using multinomial sampling, we selected *n* cells from each cell type in the mixture where *n* is 75% of the minimum number of cells available in a given cell type within the Lee el al. dataset.

### TOAST

TOAST is a recently proposed unsupervised deconvolution and feature selection algorithm which iteratively searches for cell type-specific features and performs composition estimation.(3) We use the TOAST Bioconductor package version 1.0.0 using the default 1,000 features for deconvolution. Additionally, we use NMF with KL divergence function as the deconvolution engine of TOAST.

### 3CDE

3CDE is a matrix-based deconvolution approach for bulk Hi-C data which infers non-overlapping domains of chromatin activity in each cell type from data and uses a linear combination of binary interaction information at these domains to deconvolve the contact frequency matrix.(23) We downloaded software from their Github page (https://github.com/emresefer/3cde), and ran *3cdefrac.py* with default settings. We found that the results were not usable when deconvolving multiple samples with the same underlying cell types without additional feature matching algorithms (see **Supplementary Figure 1**).

#### Simulating Bulk Hi-C Data

##### Ramani et al. Dataset

Cellular indices were downloaded from GSE84920 which included 6 libraries: ML1, ML2, ML3, ML4, PL1 and PL2.(10) For our simulations, we use data from all libraries except ML4. These libraries are composed of scHi-C data from five distinct human and mouse cell lines. Within each cell, we follow the same preprocessing procedure as outlined in Ramani *et al*. Specifically, cellular indices with fewer than 1000 unique reads, a *cis*:*trans* ratio less than 1, and cells with less than 95% of reads aligning uniquely to either the mouse or human genomes are filtered out before analysis. Additionally, we remove reads whose genomic distance was <15Kb due to self-ligation, and only considered unique reads. For the four libraries containing HAP1 and HeLa cells (ML1, ML2, PL1 and PL2), cellular indices were discarded where the proportion of sites where the non-reference allele was found was between 57% and 99%.

To account for varying levels of single-cell sequencing depth across libraries, we consider only cells with filtered reads greater than the 20^th^ quantile and less than the 90^th^ quantile of reads and across all libraries and cell types considered in the simulated mixture sample. We then downsample each cell via multinomial sampling to the number of contacts in the cell with the fewest number of contacts across all cell types considered in the sample. We construct contact matrices on the filtered and downsampled scHi-C data at three levels of data representation at 10Mb bin-pair resolution: interchromosomal contacts only, intrachromosomal contacts only, and both interchromosomal contacts and intrachromosomal contacts together. The total number of cells in each mixture sample is equal to the smallest number of cells present in a cell line after the filtering step across cells in the mixture sample.

To test proposed feature selection methods for THUNDER, we generate three cell type mixtures of GM12878, HAP1, and HeLa cells. We generate 5 replications of 12 bulk samples (3 pure samples and 9 mixture samples) which are mixtures of the three cell lines at the proportions given in **Supplementary Table 4**. These proportions are a subset of those used by Shen-Orr and Tibsherani in their simulated mixture data.(1)

##### Lee et al. Dataset

4,238 scHi-C profiles from the prefrontal cortex region of two postmortem adult human brains were downloaded from GSE130711. Non-neuronal cell types were previously identified via clustering based on CG methylation signature, followed by fine clustering of neuronal subtypes using non-CG methylation. For each cell, we removed reads with genomic distance <15kb and only considered unique reads.

We generate 5 replications of 18 mixtures of scHi-C data at 10Mb resolution consisting of 6 cell groups: oligodendrocyte (ODC), oligodendrocyte progenitor cell (OPC), astrocyte (Astro), microglia (MG), endothelial (Endo), and the 8 neuronal subtypes as one group (Neuron). Mixtures were generated at the same three resolutions of Hi-C data as the mixtures from Ramani *et al* (Supplemental Table 5).

In order to assess the robustness of the reference-based deconvolution method compared to reference-free deconvolution approaches, MuSiC, we estimated cell type proportions under three scenarios.(6) First, we estimated cell type proportions where all cell types in the mixture were present in the reference panel. Second, we randomly removed one or two cells, respectively, from the reference panel and estimated the cell type proportions of the remaining cells.

##### Window Size

In large part, the 10Mb window choice was limited by the library size of current scHi-C datasets and sparsity of contacts from which to generate synthetic bulk Hi-C datasets such that the true cell type proportions are known. Additionally, we report from our computation test on 10Kb resolution Hi-C data that THUNDER scales up to the much larger feature space of finer resolution Hi-C data. As single-cell technologies improve and with more data accumulating, we will be able to test Hi-C deconvolution methods at finer data resolutions where truth is known.

##### Feature Selection

The eleven feature selection methods either performed feature selection on the bulk Hi-C contact frequencies or on the derived cell-type specific profiles after an initial NMF fit. Strategies in the former group identify bin-pairs with high Fano Factor estimates across all samples. Strategies in the latter group identify informative bin-pairs with high cell-type-specificity and/or high variation across inferred cell types. Cell type specificity is measured by feature score within a bin-pair and across estimated cell types. Across-cell-type variation is measured by standard deviation within a bin-pair and across estimated cell types. For both metrics, we use empirical thresholds based on the distribution of these estimates across all bin-pairs for feature selection.

#### Real Data Analysis

##### Sullivan Lab eHi-C data

Anterior temporal cortex was dissected from postmortem samples from three adults of European ancestry with no known psychiatric or neurological disorder. Protocol for generating Hi-C data on these samples has been described previously(26). We applied THUNDER to the three adult samples at 1Mb resolution to match the resolution of our real-data based simulations. We ran THUNDER on intrachromosomal contacts only, and performed feature selection on each chromosome separately. To obtain the final estimated cell type proportions, we concatenated selected features across all chromosomes before running step 2 of the THUNDER algorithm. We assumed a range of possible values for the number of cells in the mixture (k = 3,…,7), and ran THUNDER for 100 iterations for both feature selection and cell type proportion estimation.

After running THUNDER, we identified bin-pairs that demonstrated specificity to each inferred cell-type-profile. Informative bin-pairs were selected as specific to each inferred cell-type-profile if the row-normalized element of the basis matrix was greater than or equal to 0.3. This threshold was chosen to select a sufficient number of bin-pairs for each feature. We then compared the unique bins in these bin-pairs with cell-type specific epigenomic annotations (described below). We assigned cell types to the THUNDER inferred cluster-specific contact profiles based on the enrichment of epigenetic features within the THUNDER bins based on the results of a chi-squared test. Finally, we compared the THUNDER estimated cell-type proportions for each labelled cluster with the distribution of cell types within cortex tissue.

##### Enhancer Annotations

We obtained cell-type-specific enhancer annotations for neurons, microglia, oligodendrocytes, and astrocytes generated from Nott *et al.* They performed ATAC-seq as well as H3K27ac and H3K4me3 chromatin immunoprecipitation sequencing on cell-type-specific nuclei. We did not consider cell-type-specific enrichments for promoters due to previous evidence supporting that promoters are mostly conserved across cell types.(34)

##### Cell Type Specifically Expressed Genes

We used cell-type-specific RNA-seq data in neurons, microglia, oligodendrocytes, and astrocytes generated by Zhang *et al.* to identify cell type specific genes.(35) We defined a cell type specific gene as a gene where the difference between the cell type specific expression and the mean expression level of all other genes was greater than one. To examine overlap with Hi-C bins, we check the region within 2kb of the gene transcription start site.

##### High-confidence regulatory chromatin interactions

High confidence regulatory chromatin interactions (HCRCIs) are genomic regions physically proximal in the nuclear 3D space. HCRCIs were identified for the three adult cortex tissue samples as described above in a previous study.(26) HCRCIs are interacts that demonstrated significant evidence of increased interaction frequency (p < 2.31 10^−11^) and overlapped with open chromatin, active histone marks, or transcription start sites of brain-expressed genes. Data were generated with two 10 Kb anchors that are ≥20 Kb and ≤2 Mb apart.

#### Computation Test with 10Kb Hi-C data

In order to assess the computational costs of THUNDER on genome-wide Hi-C data, we apply THUNDER to intrachromosomal Hi-C data at 10Kb resolution in YRI samples.(8) We randomly select 5 samples to be included in the analyses. First, we perform feature selection for each chromosome through simple parallelization. Then, we concatenate the selected features across all chromosomes for the final deconvolution estimate. We use computing time and memory usage to assess the computational efficiency for both feature selection and estimation of cell type proportions across the three datasets.

## Notes

### Competing Interest Statement

The authors have declared no competing interest.

### Summary of Updates

New simulation and real-data results with 6 cell type mixtures and an application to real Hi-C data.

