## Supplemental Information for "THUNDER: A reference-free deconvolution method to infer cell type proportions from bulk Hi-C data"

### Supplemental Note on THUNDER Feature Selection

Let $W_1(i,j)$ denote the element in the $i^{th}$ row and $j^{th}$ column of the cell type profile matrix $W_1$. Then $i = 1, \dots, p$ indicates bin-pair $i$. Let $S_{intra}$ denote the set of all intrachromosomal bin-pairs. The derivation below is for intrachromosomal bin-pairs, but the feature selection algorithm is the same for interchromosomal variants. Standard deviation across cell types for base-pair $i$ is defined as,

$$SD_i = \frac{1}{k-1}\sum_{j=1}^{k}\left(W(i,j) - \frac{1}{k}W(i,\cdot)\right)^2$$

Larger values of the standard deviation across cell types for a given bin-pair indicate greater variation in the estimated cell type specific chromatin activity.

Feature score across cell types for base-pair $i$ is defined as follows.

$$(Feature\ Score)_i = 1 + 1/log_2(k)\sum_{j=1}^{k} p(i,j)log_2\big(p(i,j)\big)$$

where $p(i,\Omega)$ is the probability that the $i$-th pairwise bin contributes to cell type $\Omega$ , i.e. $p(i,\Omega) = \frac{W_1(i,\Omega)}{\sum_{j=1}^{k} W_1(i,j)}$. Feature scores range from [0,1] with higher scores representing bin-pairs with higher cell type specificity. We find that a combination of standard deviation and feature score computed across cell types more effectively identifies informative bin-pairs than either summary statistic working individually (see **Results**).

Consider,

$$\hat{\mu}_{SD,intra} = \frac{1}{|S_{intra}|}\sum_{\{i:\, i \in S_{intra}\}} SD_i$$

$$\hat{\mu}_{FS,intra} = \frac{1}{|S_{intra}|}\sum_{\{i:\, i \in S_{intra}\}} FS_i$$

$$\hat{\sigma}_{SD,intra} = \frac{1}{|S_{intra}| - 1}\sum_{\{i:\, i \in S_{intra}\}} \left(SD_i - \hat{\mu}_{sd,intra}\right)^2$$

$$\hat{\sigma}_{FS,intra} = \frac{1}{|S_{intra}| - 1}\sum_{\{i:\, i \in S_{intra}\}} \left(FS_i - \hat{\mu}_{sd,intra}\right)^2$$

Let $FF_i = \frac{\mu_i}{\sigma_i}$ be the Fano Factor for the $i^{th}$ row of the mixture matrix. Let $e_i$ denote the row-wise maximum for for the $i^{th}$row of the cell type profile matrix. Let $\widehat{m}_{CTP}$ denote the median value of all elements of the CTP matrix. In the following table the *intra* subscript is dropped for clarity.

| Feauture Selection Method | Method definition |
|---|---|
| CTS or ICV | $FS_i > \hat{\mu}_{fs} + 3\hat{\sigma}_{fs}$ OR $SD_i > \hat{\mu}_{sd} + 3\hat{\sigma}_{sd}$ |
| CTS or ICV - Median | $FS_i > \widehat{m}_{fs} + 3\hat{s}_{fs}$ OR $SD_i > \widehat{m}_{sd} + 3\hat{s}_{sd}$ |
| CTS and ICV | $FS_i > \hat{\mu}_{fs} + 3\hat{\sigma}_{fs}$ AND $SD_i > \hat{\mu}_{sd} + 3\hat{\sigma}_{sd}$ |
| CTS and ICV - Median | $FS_i > \widehat{m}_{fs} + 3\hat{s}_{fs}$ AND $SD_i > \widehat{m}_{sd} + 3\hat{s}_{sd}$ |
| CTS | $FS_i > \hat{\mu}_{fs} + 3\hat{\sigma}_{fs}$ |
| ICV (THUNDER – inter) | $SD_i > \hat{\mu}_{sd} + 3\hat{\sigma}_{sd}$ |
| CTS – Median (THUNDER -intra) | $FS_i > \widehat{m}_{fs} + 3\hat{s}_{fs}$ |
| ICV - Median | $SD_i > \widehat{m}_{sd} + 3\hat{s}_{sd}$ |
| Top 1000 FF | Select top 1000 rows on $FF_i$ |
| Top 100 FF | Select top 100 rows on $FF_i$ |
| Kim-Park | $FS_i > \hat{\mu}_{fs} + 3\hat{\sigma}_{fs}$ AND $e_i > \widehat{m}_{CTP}$ |

Supplemental Tables 2-6

| Chr | Duration (h) | Memory (GB) | Bin-Pairs | Selected Bin-Pairs |
|---|---|---|---|---|
| **THUNDER Step 1** | | | | |
| chr1 | 4.7 | 85.1 | 2,898,116 | 57,611 |
| chr2 | 3.7 | 96.7 | 3,211,342 | 63,152 |
| chr3 | 4.5 | 70.3 | 2,742,971 | 54,380 |
| chr4 | 4.4 | 73.1 | 2,653,508 | 52,331 |
| chr5 | 4.0 | 73.1 | 2,457,251 | 48,239 |
| chr6 | 3.8 | 54.6 | 2,334,790 | 46,153 |
| chr7 | 2.5 | 52.1 | 1,972,510 | 39,288 |
| chr8 | 2.3 | 51.9 | 1,926,609 | 38,021 |
| chr9 | 1.5 | 44.0 | 1,443,866 | 28,720 |
| chr10 | 3.2 | 48.5 | 1,681,822 | 33,231 |
| chr11 | 3.7 | 51.5 | 1,727,598 | 34,530 |
| chr12 | 3.6 | 51.7 | 1,752,414 | 35,044 |
| chr13 | 2.5 | 37.6 | 1,364,537 | 26,453 |
| chr14 | 1.3 | 41.2 | 1,170,924 | 23,367 |
| chr15 | 1.0 | 31.0 | 996,396 | 20,132 |
| chr16 | 1.5 | 29.3 | 832,338 | 16,141 |
| chr17 | 1.8 | 29.6 | 841,985 | 17,422 |
| chr18 | 0.9 | 32.2 | 1,025,512 | 20,399 |
| chr19 | 0.9 | 22.1 | 468,526 | 9,114 |
| chr20 | 0.9 | 26.4 | 730,999 | 14,688 |
| chr21 | 0.7 | 22.8 | 441,879 | 8,468 |
| chr22 | 0.3 | 19.5 | 339,585 | 6,887 |
| **THUNDER Step 2** | | | | |
| Genome-Wide | 0.7 | 16.2 | 693,771 | NA |

**Supplementary Table 2**. Computational Performance on 3 YRI Samples of 10Kb Resolution Hi-C Data.

| Chr | Duration (h) | Memory (GB) | Bin-Pairs | Selected Bin-Pairs |
|---|---|---|---|---|
| **THUNDER Step 1** | | | | |
| chr1 | 5.4 | 97.3 | 3,174,333 | 65,265 |
| chr2 | 7.2 | 103.2 | 3,507,051 | 68,506 |
| chr3 | 6.3 | 92.4 | 2,988,427 | 61,700 |
| chr4 | 5.1 | 89.1 | 2,893,327 | 59,152 |
| chr5 | 4.0 | 85.7 | 2,677,352 | 53,739 |
| chr6 | 6.9 | 81.2 | 2,545,972 | 50,167 |
| chr7 | 6.1 | 69.6 | 2,163,857 | 44,196 |
| chr8 | 3.3 | 59.1 | 2,105,779 | 41,971 |
| chr9 | 2.5 | 49.0 | 1,580,008 | 32,001 |
| chr10 | 3.2 | 59.4 | 1,848,124 | 36,801 |
| chr11 | 3.1 | 57.5 | 1,896,637 | 37,615 |
| chr12 | 2.9 | 62.2 | 1,931,846 | 38,602 |
| chr13 | 2.3 | 49.2 | 1,483,383 | 29,353 |
| chr14 | 1.9 | 45.3 | 1,278,870 | 25,480 |
| chr15 | 2.1 | 35.7 | 1,094,740 | 22,468 |
| chr16 | 2.4 | 34.5 | 924,458 | 17,667 |
| chr17 | 3.8 | 37.3 | 935,763 | 19,203 |
| chr18 | 1.5 | 42.7 | 1,120,797 | 21,741 |
| chr19 | 1.5 | 23.5 | 524,543 | 9,540 |
| chr20 | 1.3 | 30.0 | 802,176 | 15,743 |
| chr21 | 1.1 | 25.2 | 485,244 | 9,076 |
| chr22 | 0.6 | 21.1 | 380,611 | 7,713 |
| **THUNDER Step 2** | | | | |
| Genome-Wide | 2.5 | 18.2 | 767,700 | NA |

**Supplementary Table 3.** Computational Performance on 5 YRI Samples of 10Kb Resolution Hi-C Data.

| Chr | Duration (h) | Memory (GB) | Bin-Pairs | Selected Bin-Pairs |
|---|---|---|---|---|
| **THUNDER Step 1** | | | | |
| chr1 | 12.6 | 110.5 | 3,538,635 | 68,709 |
| chr2 | 15.4 | 160.2 | 3,903,593 | 78,890 |
| chr3 | 20.3 | 103.8 | 3,306,003 | 68,256 |
| chr4 | 17.4 | 106.6 | 3,201,164 | 65,000 |
| chr5 | 11.0 | 114.4 | 2,967,778 | 58,602 |
| chr6 | 9.8 | 86.9 | 2,818,330 | 56,838 |
| chr7 | 12.9 | 93.5 | 2,426,518 | 50,105 |
| chr8 | 11.8 | 89.6 | 2,343,805 | 46,794 |
| chr9 | 6.6 | 73.7 | 1,761,728 | 34,939 |
| chr10 | 11.4 | 80.4 | 2,070,850 | 41,651 |
| chr11 | 9.5 | 85.1 | 2,117,742 | 42,870 |
| chr12 | 7.6 | 84.6 | 2,129,312 | 41,521 |
| chr13 | 12.1 | 63.5 | 1,638,718 | 31,997 |
| chr14 | 5.2 | 62.1 | 1,431,155 | 28,821 |
| chr15 | 6.3 | 53.3 | 1,234,104 | 24,959 |
| chr16 | 5.1 | 47.6 | 1,060,098 | 20,798 |
| chr17 | 3.7 | 43.9 | 1,075,081 | 21,785 |
| chr18 | 4.8 | 54.0 | 1,245,372 | 25,415 |
| chr19 | 3.6 | 27.9 | 616,312 | 10,771 |
| chr20 | 4.7 | 39.9 | 900,160 | 18,390 |
| chr21 | 3.8 | 28.1 | 545,011 | 10,082 |
| chr22 | 1.5 | 27.1 | 440,207 | 8,700 |
| **THUNDER Step 2** | | | | |
| Genome-Wide | 4.1 | 28.6 | 855,893 | NA |

**Supplementary Table 4.** Computational Performance on 10 YRI Samples of 10Kb Resolution Hi-C Data.

Table 1: Mixing proportions for GM12878, HAP1, and HeLa mixtures.

| Sample Number | GM12878 | HAP1 | HeLa |
|---|---|---|---|
| 1 | 1.00 | 0.00 | 0.00 |
| 2 | 0.00 | 1.00 | 0.00 |
| 3 | 0.00 | 0.00 | 1.00 |
| 4 | 0.70 | 0.25 | 0.05 |
| 5 | 0.70 | 0.05 | 0.25 |
| 6 | 0.05 | 0.70 | 0.25 |
| 7 | 0.05 | 0.25 | 0.70 |
| 8 | 0.25 | 0.05 | 0.70 |
| 9 | 0.25 | 0.70 | 0.05 |
| 10 | 0.45 | 0.45 | 0.10 |
| 11 | 0.45 | 0.10 | 0.45 |
| 12 | 0.10 | 0.45 | 0.45 |

**Supplementary Table 5.** Mixing Proportions for GM12878, HAP1, and HeLa Simulations.

| Sample Number | ODC | Astro | MG | Endo | OPC | NeuronMix |
|---|---|---|---|---|---|---|
| 1 | 1.00 | 0.00 | 0.00 | 0.00 | 0.00 | 0.00 |
| 2 | 0.00 | 1.00 | 0.00 | 0.00 | 0.00 | 0.00 |
| 3 | 0.00 | 0.00 | 1.00 | 0.00 | 0.00 | 0.00 |
| 4 | 0.00 | 0.00 | 0.00 | 1.00 | 0.00 | 0.00 |
| 5 | 0.00 | 0.00 | 0.00 | 0.00 | 1.00 | 0.00 |
| 6 | 0.00 | 0.00 | 0.00 | 0.00 | 0.00 | 1.00 |
| 7 | 0.49 | 0.10 | 0.10 | 0.10 | 0.10 | 0.10 |
| 8 | 0.10 | 0.49 | 0.10 | 0.10 | 0.10 | 0.10 |
| 9 | 0.10 | 0.10 | 0.49 | 0.10 | 0.10 | 0.10 |
| 10 | 0.10 | 0.10 | 0.10 | 0.49 | 0.10 | 0.10 |
| 11 | 0.10 | 0.10 | 0.10 | 0.10 | 0.49 | 0.10 |
| 12 | 0.10 | 0.10 | 0.10 | 0.10 | 0.10 | 0.49 |
| 13 | 0.00 | 0.20 | 0.20 | 0.20 | 0.20 | 0.20 |
| 14 | 0.20 | 0.00 | 0.20 | 0.20 | 0.20 | 0.20 |
| 15 | 0.20 | 0.20 | 0.00 | 0.20 | 0.20 | 0.20 |
| 16 | 0.20 | 0.20 | 0.20 | 0.00 | 0.20 | 0.20 |
| 17 | 0.20 | 0.20 | 0.20 | 0.20 | 0.00 | 0.20 |
| 18 | 0.20 | 0.20 | 0.20 | 0.20 | 0.20 | 0.00 |

**Supplemental Table 6.** Mixing proportions for Lee *et al.* mixtures of 6

### Supplemental Figures

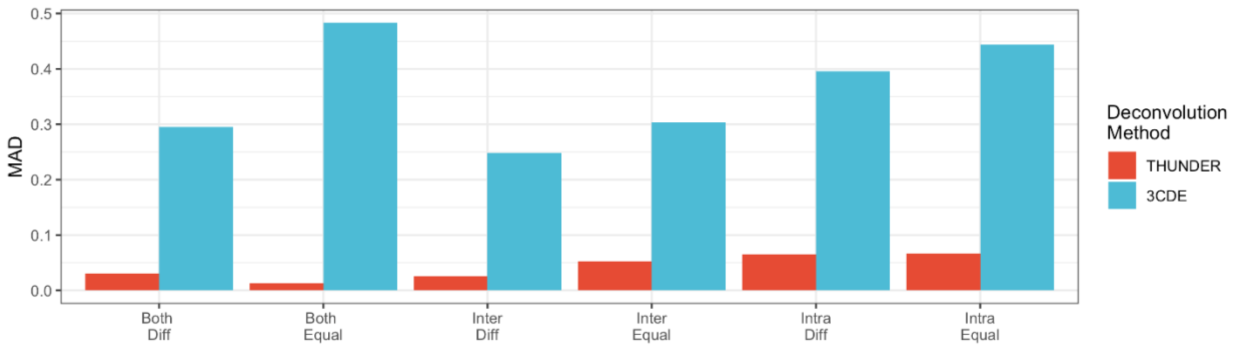


**Supplementary Figure 1.** Performance of *THUNDER* and 3CDE on HAP1 and HeLa Simulated Mixtures**.** We see that in several simulations, 3CDE achieves near the maximum mean absolute deviation from true cell type proportions (0.5).We do not test 3CDE in further simulations because of its inability to handle multiple Hi-C samples simultaneously.

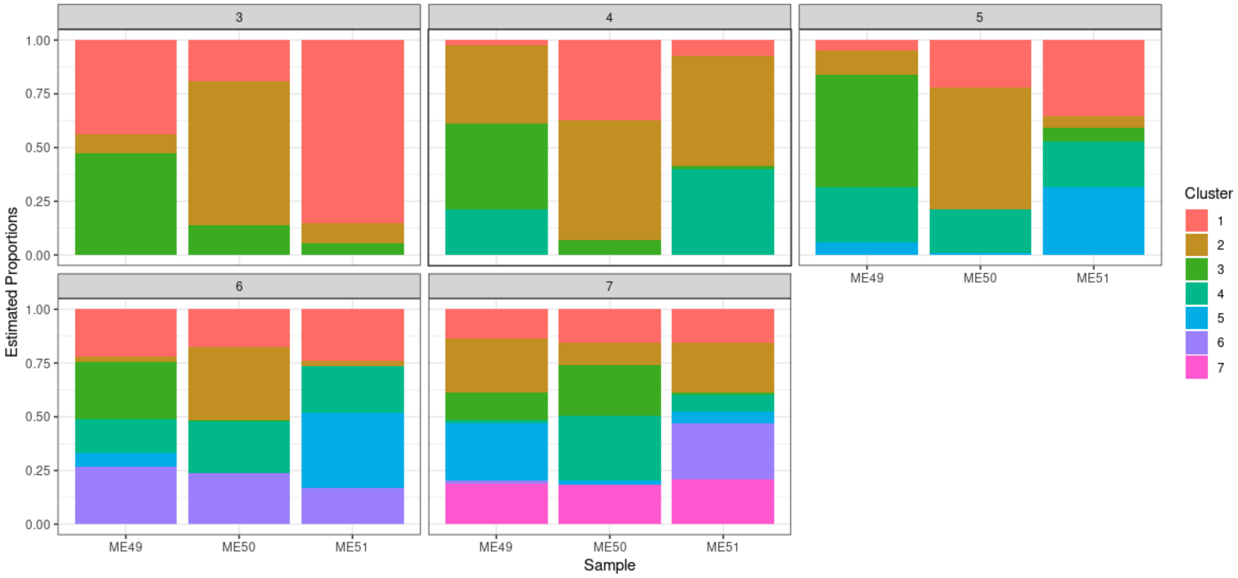


**Supplemental Figure 2.** THUNDER cell type proportion estimates across a range of k=3,..,7 values in NMF deconvolution for deconvolution of Giusti-Rodriguez *et al.* Hi-C data.
